# Structural compatibility enables functional co-assembly of Kir2.1 channels and the voltage-sensing domain of voltage-sensitive phosphatases

**DOI:** 10.64898/2026.08.07.743441

**Authors:** Anagha Gopalakrishnan Nair, Philipp Rühl, Lina Seeber, Toshinori Hoshi, Roland Schönherr, Stefan H. Heinemann

## Abstract

Voltage-sensing domains (VSDs), which are integral parts of voltage-gated K^+^ channel (K_V_) proteins, are highly modular protein components that function largely independently of ion-conducting pores – a property exploited in genetically encoded voltage indicators (GEVIs). Conversely, inward-rectifier K_+_ channels such as Kir2.1 possess a pore-only architecture and lack a VSD. Here, we demonstrate an unexpected and functionally relevant structural compatibility between the independently evolved pore-only and VSD-only membrane protein families. When co-expressed, Kir2.1 and ASAP-type GEVIs form complexes that constrain VSD movement and markedly interfere with voltage-dependent fluorescence responses. Molecular modeling combined with targeted mutagenesis identified a conserved hydrophobic interface that mediates this interaction. A single bulky substitution in the VSD of the GEVI rEstus-NI (A79W) disrupted the impact of Kir2.1 while preserving the GEVI’s voltage-sensing performance. These findings suggest that pore-only and VSD-only proteins can assemble into functional K_V_-like architectures, and highlight that membrane proteins may engage in unexpected interactions capable of altering experimental readouts in physiological voltage imaging studies. The study also raises the possibility that independently functional membrane proteins may assemble into previously unrecognized higher-order complexes with distinct functional properties under native physiological conditions.

## Introduction

Potassium (K⁺) channels regulate membrane excitability and resting membrane potential in virtually every cell type. Functional K⁺ channels are composed of pore-forming α subunits, which determine ion selectivity, the mode of activation (gating), and the overall channel protein structure. The α subunits of the evolutionarily oldest types of K⁺ channels consist of only two transmembrane segments (2TM) connected by a pore loop (Fig. 1*A*), thus forming a pore domain (PD). Four α subunits assemble to a largely voltage-independent K⁺ conducting channel [1, 2]. This architecture is found in most prokaryotic channels and is preserved in inward-rectifying K⁺ (Kir) channels in mammals. It is believed that duplication of 2TM subunits [3] has given rise to 4TM K⁺ channels (two-pore channels, K2P), such as TWIK, TREK, and TASK.

**Figure 1.**
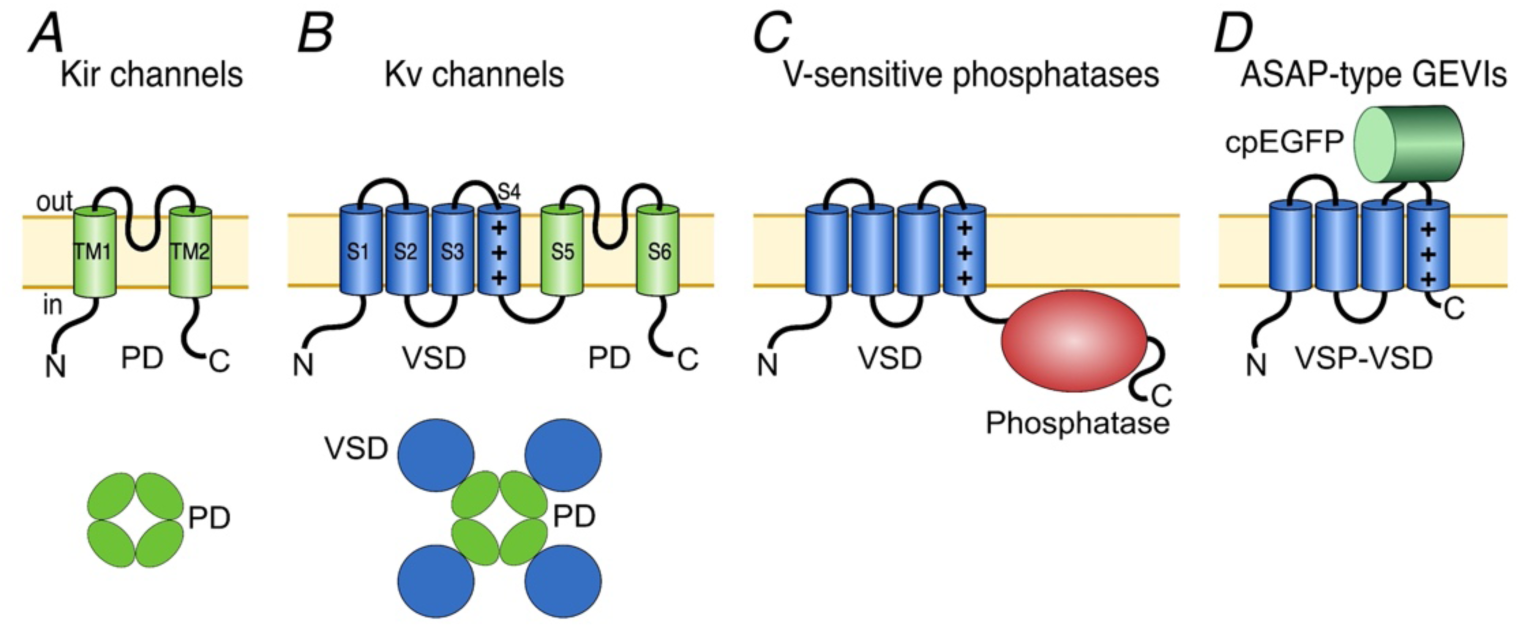
Structural organization of VSD-based sensors and K^+^ channel architectures. *A*, Membrane topology of a 2TM subunit of inward-rectifier K⁺ (Kir) channels. Kir channel subunits are composed of two transmembrane helices (TM1 and TM2) and a pore loop (*top*). Four Kir subunits assemble into a functional tetrameric channel (*bottom*). *B*, Membrane topology of an α subunit of 6TM channels, such as voltage-gated K^+^ (KV) channels, with a voltage-sensing domain (VSD, blue) formed by transmembrane segments S1-S4 and a pore domain (PD, green) formed by transmembrane segments S5 and S6 with the intervening pore loop that constitutes the ion-selective pore (*top*). The number of positive charges in S4 shown is just for illustrative purposes. Four KV channel subunits assemble to form a functional tetrameric channel (*bottom*). *C*, Membrane topology of a voltage-sensing phosphatase (VSP), consisting of a VSD coupled to a cytoplasmic phosphatase domain (red). *D*, Topology of a VSD-based genetically encoded voltage indicator (GEVI) of the ASAP family, in which a circularly permuted EGFP (cpEGFP, green) is inserted between S3 and S4 of the VSD. VSPs as well ASAP-type GEVIs may also exist as dimers.

In contrast, α subunits of the voltage-gated K^+^ (K_V_) channel consist of six transmembrane segments (6TM). The first four segments (S1-S4) constitute the voltage-sensing domain (VSD), while the remaining segments share structural similarity with 2TM channels and form the PD (Fig. 1*B*). S4 of the VSD typically contains several positively charged amino acids that respond to changes in the transmembrane voltage (*V*_m_), thus leading to conformational changes that ultimately open or close the K_+_-selective pore [4].

VSDs are highly modular structures capable of conferring voltage sensitivity to other protein domains [5]. As modular protein domains, VSDs are found in a variety of other voltage-dependent ion channels, such as voltage-gated sodium (Na_V_) and calcium (Ca_V_) channels, and can even function independently of a pore domain, as seen in voltage-gated proton channels (HV) [5–8]. This profound modularity is further exemplified by voltage-sensitive phosphatases (VSPs) such as *TPTE2*, in which a VSD is coupled to a PTEN-like (phosphatase and tensin homolog) domain to mediate the voltage-dependent dephosphorylation of membrane phospholipids such as PI(3,4)P₂, PI(4,5)P₂, and PI(3,4,5)P₃ (Fig. 1*C*) [9, 10].

This modular architecture has been successfully exploited to generate genetically encoded voltage indicators (GEVIs), in which a fluorescent protein is fused to a highly optimized VSD to report transmembrane voltage changes optically (Fig. 1*D*). Among these, GEVIs of the ASAP type (Accelerated Sensor(s) of Action Potentials), based on the VSD of the *Gallus gallus* VSP (gVSP), are widely used in physiological studies due to their high brightness, sensitivity, and fast kinetics. These sensors contain a circularly permuted enhanced GFP (cpEGFP) inserted between the VSD’s S3 and S4 helices, which alters its fluorescence in response to VSD conformational changes [11].

Typically, VSDs and their respective effector domains, whether a PD, a PTEN domain, or a fluorophore, are covalently linked within a polypeptide. However, experimental separation of the VSD from the PDs of ether-à go-go-type α subunits (“split channels”) has revealed that these domains can co-assemble into functional K_V_ channels without covalent bonds linking the VSD to the PD [12, 13]. Because these domains co-evolved within the same protein, their structural compatibility is expected. Whether a 2TM protein, such as in Kir channels, can form functional complexes with a VSD from a different protein family, such as voltage-sensitive phosphatases, remains unknown.

Inwardly rectifying Kir channels typically stabilize the resting membrane potential. Their α subunits, encoded by *KCNJ* genes, can form homo- or heteromeric channel complexes [14, 15] (Fig. 1*A*). Among these, Kir2.1 (*KCNJ2*) mediates the inward-rectifier K^+^ current (I_K1_), which contributes to resting *V*_m_ maintenance and late-phase repolarization in the heart [16, 17]. Kir2.1 channels are also present in skeletal muscle [18], neurons [19], and glial cells [20].

Here, we report that Kir2.1 channel α subunits specifically alter the voltage-dependent optical properties of ASAP-type GEVIs, possibly by forming a non-covalent VSD-2TM complex. Using structure-guided mutagenesis and whole-cell patch-clamp fluorometry, we identified the specific interaction surface between Kir2.1 and the GEVI. By introducing sterically bulky residues into this interface, we successfully disrupted this interaction. Notably, a single A79W mutation in the S1 segment of the GEVI’s VSD was sufficient to render the sensor largely insensitive to Kir2.1 co-expression. The findings reveal that GEVIs are not necessarily isolated from interactions with endogenous membrane proteins, raising potentially important implications for VSD-based physiological voltage imaging.

## Methods

### Structural prediction with AlphaFold 3

A hetero-octameric complex consisting of four rEstus [21] and four mouse Kir2.1 (mKir2.1) subunits was modeled *in silico* using the AlphaFold 3 server [22]. In addition to an overall ranking score, the predicted template modeling score (pTM) and the interface predicted template modeling score (ipTM), both ranging from 0 to 1, were determined. The resulting model was used to identify putative interface residues to guide mutagenesis. Structural models were visualized and analyzed using UCSF ChimeraX [23] and PyMOL Molecular Graphics System (Version 2.6.2 Schrödinger, LLC).

### Molecular dynamics simulation

The best-scoring structural model predicted by AlphaFold 3 was energy optimized in all-atom molecular dynamics simulation runs with CHARMM-GUI [24] and NAMD2/3 using the CHARMM36m force field. The lipid bilayer consisted of cholesterol (20%), dipalmitoylphosphatidylcholine (DPPC, 40%) and dioleoylphosphatidylcholine (DOPC, 40%) in the extracellular leaflet, as well as cholesterol (20%), DPPC (25%), DOPC (25%), dipalmitoylphosphatidylethanolamine (DPPE, 10%), dioleoylphosphatidylethanolamine (DOPE, 10%), dipalmitoylphosphatidylserine (DPPS, 5%), and dioleoylphosphatidylserine (DOPS, 5%) in the intracellular leaflet [25]. The periodic boundary simulation box, measuring approximately 15 nm x 15 nm x 23.2 nm, contained water and 150 mM KCl, totaling 499,751 atoms (Supplementary Fig. 5*A*). The system was equilibrated according to the default parameter values suggested by CHARMM-GUI, and the simulations were performed in the NPT ensemble at 37°C. A molecular dynamics simulation of 100 ns was analyzed by measuring the time course of distances between the α carbon atoms (CA) of residues of interest.

### Molecular biology

Expression plasmids of the following GEVIs were constructed using standard molecular biology methods: rEstus-NI [26], ASAP3 [27], rEstus [21], JEDI-1P [28], ASAP5 [29], and rEstus2s [30].

The following ion channel constructs were used: mouse Kir2.1 (mKir2.1, *KCNJ2*, NP_032451.1), chicken Kir2.1 (gKir2.1, *KCNJ2*, NP_990701.2), human Kir2.1 (hKir2.1, *KCNJ2*, NP_000882.1), chicken Kir2.2 (gKir2.2, *KCNJ12*, NP_001383542.1), rat Kir1.1 (rKir1.1, *KCNJ1*, NP_058719.2), human K2p9.1 (hK2p9.1, hTASK3, *KCNK9*, NP_001269463.1), human K_V_10.1 (hK_V_10.1, hEAG1, *KCNH1*, NP_002229.1), human K_V_10.2 (hK_V_10.2, hEAG2, *KCNH5*, NP_647479.2), human K_Ca_3.1 (hK_Ca_3.1*, KCNN4*, NP_002241.1), human K_Ca_1.1 (hK_Ca_1.1, hBKα*, KCNMA1*, NP_002238.2), and the auxiliary human BKβ1 subunit (Sloβ1, *KCNMB1*, NP_004128.1).

Mutants of rEstus-NI, rEstus2s, and mKir2.1 were generated by site-directed mutagenesis. All variants were cloned into the pcDNA3.1 vector. Sequences were validated by Sanger sequencing (Eurofins).

### Sequence alignments

Pairwise global sequence alignments of the TM1 and TM2 domains from human Kir2.1, Kir2.2 and Kir1.1 channels were performed using EMBOSS Needle (Needleman-Wunsch algorithm) via the EMBL-EBI Job Dispatcher web service [31], with default parameters (BLOSUM62 substitution matrix, gap open penalty = 10.0, gap extend penalty = 0.5). Transmembrane domain boundaries of TM1 and TM2 for each channel were obtained from the corresponding UniProt entry (Kir1.1: P48048; Kir2.1: P63252; Kir2.2: Q14500).

### Cell culture

Human embryonic kidney 293T cells (HEK293T; CAMR, Porton Down, Salisbury, UK) were cultured in Dulbecco’s Modified Eagle’s Medium/Nutrient Mixture F-12 (DMEM/F-12; Thermo Fisher Scientific, Waltham, MA, USA) supplemented with 10% fetal bovine serum (FBS). Cells were maintained at 37°C in a humidified incubator with 5% CO₂.

For electrophysiology and fluorescence imaging experiments, cells were seeded onto 35-mm glass-bottom dishes (Ibidi, Martinsried, Germany) at a density of 10,000 cells per dish. The following day, cells were transfected using the ROTI^®^Fect transfection kit (Carl Roth, Karlsruhe, Germany). For control experiments, 1 µg of the corresponding GEVI DNA was transfected per dish. For experiments investigating channel effects on GEVIs, cells were co-transfected with 0.5 µg GEVI DNA and 0.5 µg ion channel DNA per dish. Simultaneous electrophysiology and fluorescence imaging experiments were performed 24 h after transfection.

### Simultaneous electrophysiology and fluorescence imaging

Fluorescence imaging recordings were performed using an Axio Observer inverted microscope (Carl Zeiss, Jena, Germany). Fluorophores were excited with 480-nm and 400-nm light from an XBO lamp coupled to a Polychrome V monochromator (T.I.L.L. Photonics, Gräfelfing, Germany). The epifluorescence filter cube comprised 492/SP excitation filter, FT495 beam splitter, and 525/50 emission bandpass.

Whole-cell patch-clamp recordings were performed using an EPC10 amplifier and PatchMaster acquisition software (HEKA Elektronik, Lambrecht, Germany). Patch pipettes were fabricated from borosilicate glass capillaries with filament, coated with dental wax, and fire-polished to achieve resistances of 1-2 MΩ. Cells with series resistances above 5 MΩ were discarded. The series resistance was electronically compensated by up to 75% aiming to limit the voltage error to less than 10 mV.

Transiently transfected cells were voltage-clamped in the whole-cell configuration using voltage step protocols ranging from -120 to 80 mV. Before each 480 nm excitation pulse, cells were pre-illuminated with 400 nm light to counteract photoswitching induced by 480 nm excitation [21]. GEVI fluorescence images were recorded using a ProgRes MFcool camera (JenOptik, Jena, Germany), controlled by the PatchMaster/SmartLux software combination (HEKA Elektronik). Images were acquired with an exposure time of 100 ms using a 40× oil immersion objective. A typical pulse protocol is depicted in Fig. 2*B*.

**Figure 2.**
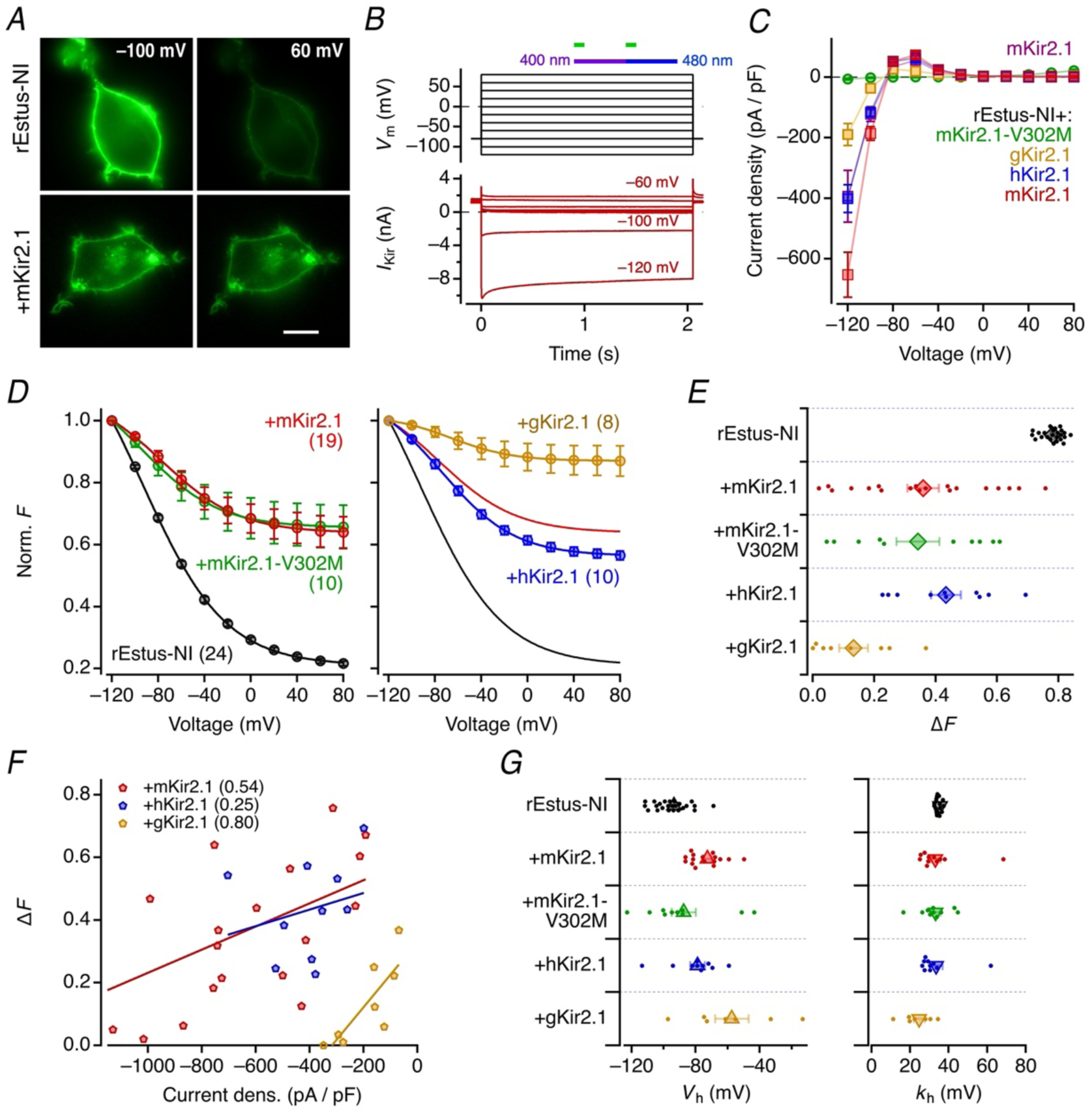
Kir2.1 co-expression impairs the voltage sensitivity of rEstus-NI. *A*, Fluorescence images of HEK293T cells expressing rEstus-NI alone (*top*) or co-expressed with mouse Kir2.1 (+mKir2.1, *bottom*) voltage-clamped at -100 mV and 60 mV. Co-expression of mKir2.1 visibly attenuates the voltage-dependent fluorescence change of rEstus-NI. Scale bar, 10 µm. *B*, Experimental protocol (*top*) and representative current recordings from a HEK293T cell expressing rEstus-NI along with mKir2.1 (*bottom*). The horizontal bars in the protocol indicate stimulation with 400- and 480-nm light as well as camera triggers (green). Images taken during 480-nm illumination were used for further analysis. Pre-illumination at 400 nm recovered part of the sensor from photoswitching. *C*, Mean whole-cell current density–voltage (*I*–*V*) relationships of mKir2.1 expressed alone (magenta) or rEstus-NI co-expressed with the indicated Kir2.1 channel types. The mKir2.1-V302M mutant (green) does not produce current. *D*, Fluorescence–voltage (*F*–*V*) relationships of rEstus-NI recorded under whole-cell voltage clamp, normalized to the fluorescence at -120 mV. Data are from rEstus-NI alone (black), co-expressed with mKir2.1 (red), or co-expressed with mKir2.1-V302M (green). Solid curves indicate Boltzmann fits (Eq. 1). Data are means ± SEM; the numbers of cells (*n*) are indicated in parentheses. *right*: As in the left panel for rEstus-NI with human Kir2.1 (hKir2.1) or chicken Kir2.1 (gKir2.1); for reference, results for rEstus-NI and rEstus-NI co-expressed with mKir2.1 are shown as black and red curves, respectively. *E*, Change in relative fluorescence of rEstus-NI between -120 and 80 mV derived from fits to individual cell data as in (*D*). *F*, Correlation of Δ*F* of rEstus-NI and current density at -120 mV of the indicated co-expression Kir2.1 channel isoforms. Regression lines are superimposed for clarity; correlation coefficients (*r*) are given in parentheses. *G*, Mean half-maximal voltage of rEstus-NI fluorescence (*V*h, *left*) and the corresponding slope factor (*k*h, *right*) from data fits in (*D*). Data in (*E*) and (*G*) are means ± SEM; individual data points are indicated as dots.

The extracellular bath solution contained (in mM): 146 NaCl, 4 KCl, 2 CaCl₂, 2 MgCl₂, and 10 HEPES, adjusted to pH 7.4 with NaOH. The intracellular solution contained (in mM): 130 KCl, 2.5 MgCl₂, 10 EGTA, and 10 HEPES, adjusted to pH 7.4 with KOH. For experiments involving expression of hK_V_10.1 channels, the intracellular solution contained (in mM) 130 CsCl, 1 MgCl₂, 10 HEPES, and 10 EGTA, adjusted to pH 7.4 with CsOH, to limit the outward current. For the same reason, currents through hK2_P_9.1 channels were measured in an intracellular solution with reduced K_+_ concentration (in mM): 14 KCl, 116 NaCl, 2.5 MgCl₂, 10 HEPES, and 10 EGTA, adjusted to pH 7.4 with KOH. All recordings were performed at room temperature (20-23°C).

For excitation-spectrum measurements, the culture medium was replaced with 2 ml of extracellular solution supplemented with 5 mM glucose. Cells were incubated in this solution for 15 min at room temperature to allow stabilization of the resting membrane potential. Fluorescence imaging was performed using the microscope, filter set, and camera settings described above. Images were acquired at each wavelength with an exposure time of 200 ms. The measured fluorescence intensities were corrected for the wavelength-dependent intensity of the excitation light. The resulting excitation spectra were subsequently normalized to the fluorescence intensity measured at 470 nm.

### Data analysis

Raw fluorescence images were analyzed by manually selecting regions of interest (ROIs) encompassing plasma membrane areas. Background fluorescence was subtracted. The resulting mean fluorescence intensity was normalized to values obtained at -120 mV to yield *F*. The dependence of *F* on *V*ₘ was described with a Boltzmann-type equation

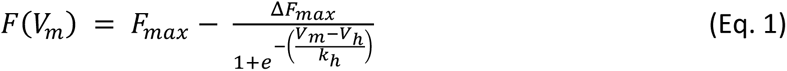

with the maximum fluorescence (*F*_max_), the maximal change in fluorescence (Δ*F*_max_), the voltage of half-maximal brightness change (*V*_h_), and the slope factor describing the steepness of the response (*k*_h_). From these parameters, the change in normalized fluorescence in the experimentally accessed voltage range between -120 and 80 mV (Δ*F*) was calculated. A similar approach was followed to describe the voltage dependence of the fluorescence ratio obtained at 480 and 400 nm excitation: *F*_480_ / *F*_400_ (*r*), where in Eq. 1 *r*_max_ and Δ*r*_max_ replace *F*_max_ and Δ*F*_max_, respectively.

To characterize fluorescence kinetics upon *V*_m_ changes, cells were subjected to a voltage step from -80 to 40 mV with a pulse duration of 512 ms. Changes in *F* following depolarization and repolarization were analyzed to assess differences in ON and OFF kinetics between the rEstus2s and rEstus2s-A79W constructs by calculating the difference in the maximum fluorescence obtained at 40 mV.

Ion currents were measured by taking the mean of the second half of the 2-s pulses. Wherever possible, linear leak was corrected offline using FITMASTER NEXT (Multi Channel Systems MCS GmbH, Reutlingen, Germany). Fluorescence images were analyzed using Fiji software [32], and final data analysis and figure generation were performed using Igor Pro 9 software (WaveMetrics, Lake Oswego, OR, USA).

Average data are presented as means ± SEM with sample sizes (*n*) and all individual data points shown as dots. *p* values, based on unpaired Student’s tests, are provided as operational data descriptors. A post-hoc Bonferroni correction was applied for multiple comparisons.

## Results

### Kir2.1 channels impair the voltage response of ASAP-type GEVIs

Kir2.1 channels are routinely used to hyperpolarize HEK293T cells during development of new GEVIs (e.g.,[27–29, 33]). We observed that coexpression of the ASAP-type GEVI rEstus-NI [26] in HEK293T cells with mouse (*Mus musculus*) Kir2.1 (mKir2.1), caused a substantial reduction in the fluorescence change between -100 and 60 mV, whereas membrane localization was not noticeably altered (Fig. 2*A*). With mKir2.1 expressed, application of a series of voltage steps ranging from -120 to 80 mV (Fig. 2*B*) revealed typical inwardly rectifying K_+_ currents irrespective of the presence of rEstus-NI (Fig. 2*B*, *C*).

The voltage dependence of rEstus-NI fluorescence (Fig. 2*D*, *left*) was described with a Boltzmann distribution (Eq. 1) and yielded a relative change in fluorescence between -120 and 80 mV (Δ*F*) of 0.78 ± 0.01 (*n* = 24) under control conditions but only 0.36 ± 0.05 (*n* = 19) in the presence of mKir2.1. To examine whether the apparent influence of mKir2.1 on rEstus-NI is related to the K_+_ current it mediates, we also measured the non-conducting mutant mKir2.1-V302M [34]. mKir2.1-V302M did not produce inwardly rectifying K_+_ current (Fig. 2*C*) but it markedly diminished Δ*F* of rEstus-NI to 0.34 ± 0.07 (*n* = 10; Fig. 2*D*, *left*).

Since the VSD of rEstus-NI originates from chicken (*Gallus gallus*) VSP (gVSP), we examined the effect of chicken Kir2.1 (gKir2.1) on rEstus-NI. This channel also mediated inwardly rectifying K_+_ currents in HEK293T cells (Fig. 2*C*), and, when co-expressed with rEstus-NI, it noticeably diminished the GEVI’s voltage sensitivity: Δ*F* of 0.13 ± 0.05 (*n* = 8; Fig. 2*D*, *right*). Human Kir2.1 (hKir2.1) also markedly diminished Δ*F* of rEstus-NI to 0.43 ± 0.05 (*n* = 10) (Fig. 2*D*, *right*). As shown in Fig. 2*E* for all individual experiments, the scatter in Δ*F* is relatively small for rEstus-NI alone but strongly increased in the presence of any of the Kir2.1 isoforms; occasionally, co-expression of mKir2.1 or gKir2.1 resulted in a complete lack of voltage dependence in rEstus-NI fluorescence (Δ*F* close to zero). This is likely explained by the weak control of the expression ratio between mKir2.1 and rEstus-NI, which is supported by a positive correlation of Δ*F* with the current density obtained at -120 mV (Fig. 2*F*). For gKir2.1, a current density at -120 mV of about -300 pA/pF already resulted in a near-total abolition of rEstus-NI’s voltage dependence, while an equivalent impact of mKir2.1 was achieved at a current density greater than -1000 pA/pF. This also suggests a stronger influence of chicken Kir2.1 than mouse or human Kir2.1 on rEstus-NI with its chicken-based VSD.

There was a depolarizing shift in the voltage of half-maximal fluorescence change (*V*_h_) and a decrease in the slope factor (*k*_h_) (greater steepness) with Kir2.1 co-expression (Fig. 2*G*). The correlation of Δ*F* with *V*_h_ and *k*_h_ (Supplementary Fig. 1) supports the trend that *V*_h_ approaches values around -50 mV for small Δ*F* values, i.e., strong overexpression of Kir2.1, compared to -100 mV without Kir2.1 expression. There is also a trend in *k*_h_ becoming smaller with diminished Δ*F* values; under these conditions *k*_h_ determination is compromised, thus resulting in a greater scatter.

rEstus-NI was optimized for its application in fluorescence-lifetime imaging microscopy (FLIM) [26], but other ASAP-type GEVIs are more frequently used for fluorescence intensity measurements. We therefore examined whether mKir2.1 co-expression would also have a functional impact on ASAP3 [27], rEstus [21], JEDI-1P [28], ASAP5 [29], and rEstus2s [30]. We chose mKir2.1 rather than gKir2.1 because in typical GEVI applications an interference with a mammalian channel subunit is a more likely scenario. Co-expression of mKir2.1 clearly altered the *F*–*V* relationships of all tested GEVIs compared with cells expressing the sensors alone (Fig. 3*A*). However, the magnitude of this effect varied among the sensors (Fig. 3*B*). Since all sensors have different absolute changes in fluorescence between -120 and 80 mV, we related Δ*F* to the control values without mKir2.1 expression (Δ*F*_0_), yielding the following mean (Δ*F*_0_ - Δ*F*) / Δ*F*_0_ values: rEstus-NI, 0.54; ASAP3, 0.49; rEstus, 0.21; JEDI-1P, 0.19; ASAP5, 0.14; rEstus2s, 0.29. There was always a significant reduction of Δ*F* by co-expression of mKir2.1 (Fig. 3*B*); the largest impact was noted for rEstus-NI, followed by ASAP3. Analysis of the voltage dependence of GEVI fluorescence using Eq. 1 showed that all tested GEVIs exhibited the same trend as rEstus-NI, namely a shift in *V*_h_ in the depolarizing direction and, except for rEstus, a moderate decrease in *k*_h_ (Supplementary Fig. 2*A*).

**Figure 3.**
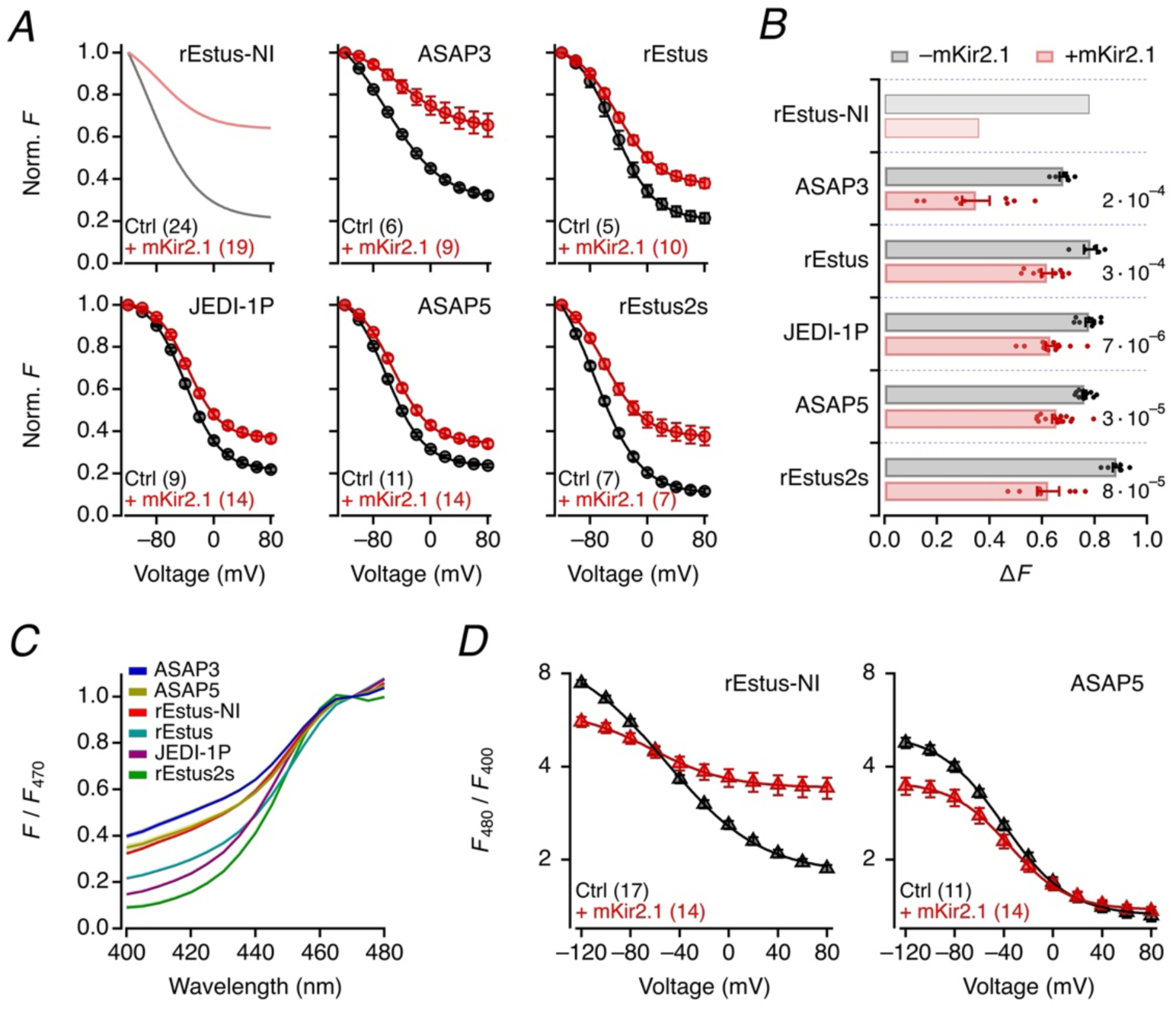
Co-expression of mKir2.1 alters the fluorescence–voltage relationship of ASAP-type GEVIs. *A*, Fluorescence–voltage (*F*–*V*) relationships recorded under whole-cell voltage clamp of HEK293T cells expressing ASAP3, rEstus, JEDI-1P, ASAP5, or rEstus2s. The fluorescence was normalized to the value at -120 mV. Data are from the respective GEVIs alone (black) or co-expressed with mKir2.1 (red). Solid curves are the results of Boltzmann fits (Eq. 1). Data are means ± SEM, *n* in parentheses. *B*, Mean relative change in fluorescence between -120 and 80 mV for the indicated GEVIs without or with co-expression of mKir2.1; mean values for rEstus-NI are shown as reference. *p* values for non-paired t-tests without and with mKir2.1 are indicated. For *V*h and *k*h values, see Supplementary Fig. 2A. *C*, Excitation spectra of HEK293T cells expressing the indicated GEVIs without voltage-clamp control (*V*m approximately -45 mV), corrected for the excitation intensity and normalized to the values obtained at 470 nm. *D*, Fluorescence ratios *F*480 / *F*400 (*r*) as a function of voltage for rEstus-NI and ASAP5 without and with co-expression of mKir2.1. The superimposed curves are results of fits according to Eq.1 yielding the following parameters: rEstus-NI (with mKir2.1), *r*max = 9.47 ± 0.23 (6.06 ± 0.09), Δ*r*max = -7.66 ± 0.25 (-2.64 ± 0.10), *V*h = -81.7 ± 2.4 mV (-71.4 ± 2.4 mV), *k*h = 36.9 ± 1.3 mV (30.8 ± 1.6 mV); ASAP5 (with mKir2.1), *r*max = 5.10 ± 0.04 (3.58 ± 0.03), Δ*r*max = -3.78 ± 0.05 (-2.20 ± 0.04), *V*h = -57.5 ± 0.7 mV (-46.7 ± 1.1 mV), *k*h = 25.6 ± 0.6 mV (23.4 ± 1.0 mV).

To better understand the mechanistic basis for the Kir2.1-induced changes in the fluorescence-voltage relationship, we examined the fluorescence ratio obtained with excitation at 480 and 400 nm (*F*_480_/*F*_400_, *r*). When the VSD is coupled to an EGFP-based fluorescent chromophore, as in GEVIs, only the deprotonated anionic chromophore is sensitive to the VSD environment as previously shown for rEstus and ASAP3 [21] and here for ASAP5 and rEstus-NI (Supplementary Fig. 2*B*). The protonation state of the chromophore can be inferred by excitation with different wavelength lights; excitation at 480 nm preferentially probes the deprotonated state, while excitation at 400 nm reports the protonated neutral state. Excitation spectra of six ASAP-type GEVIs (Fig. 3*C*) reveal that rEstus-NI, ASAP5 and ASAP3 showed strong emission with 400 nm excitation, rendering them suitable for the *F*_480_/*F*_400_ analysis to probe the fluorophore status. In addition to rEstus-NI, ASAP5 was selected for detailed analysis because it has a different linker between the cpEGFP and the VSD, compared with rEstus-NI, rEstus, ASAP3, and rEstus2s.

The voltage dependence of *F*_480_/*F*_400_ of rEstus-NI and ASAP5 by co-expression of mKir2.1 is summarized in Fig. 3*D*. With both rEstus-NI and ASAP5, the ratio was greater at hyperpolarized voltages and decreased with progressive depolarization, suggesting that a greater relative contribution of the deprotonated chromophore at negative voltages. This voltage dependence was clearly diminished by mKir2.1 co-expression albeit differentially in rEstus-NI and in ASAP5. In rEstus-NI, the voltage-dependent *r* change was compressed so that the co-expression decreased the *r* value at hyperpolarized voltages and increased it at depolarized voltages. In contrast, for ASAP5, mKir2.1 co-expression only decreased the *r* value at negative voltages. The differential effects of co-expression with mKir2.1 suggest that the linker between the cpEGFP and the VSD may influence the coupling between the VSD and the chromophore deprotonated-protonated distribution.

### Kir2.1 interferes specifically with the voltage-sensing domain of gVSP

To determine whether the observed interference of Kir2.1 with rEstus-NI is specific to Kir2.1, we examined various K⁺ channels with different structural architectures (Fig. 4). K_+_ channels can be broadly classified based on the transmembrane topology of their α subunits. Kir2.1 channels have a 2TM pore-only architecture (Fig. 1*A*). Phylogenetic analysis places Kir2.1 in close proximity to Kir2.2, while Kir1.1 forms a more distantly related branch; even restricted to the TM1/TM2 transmembrane regions, Kir2.1 is more similar to Kir2.2 than to Kir1.1 (Supplementary Fig. 3).

**Figure 4.**
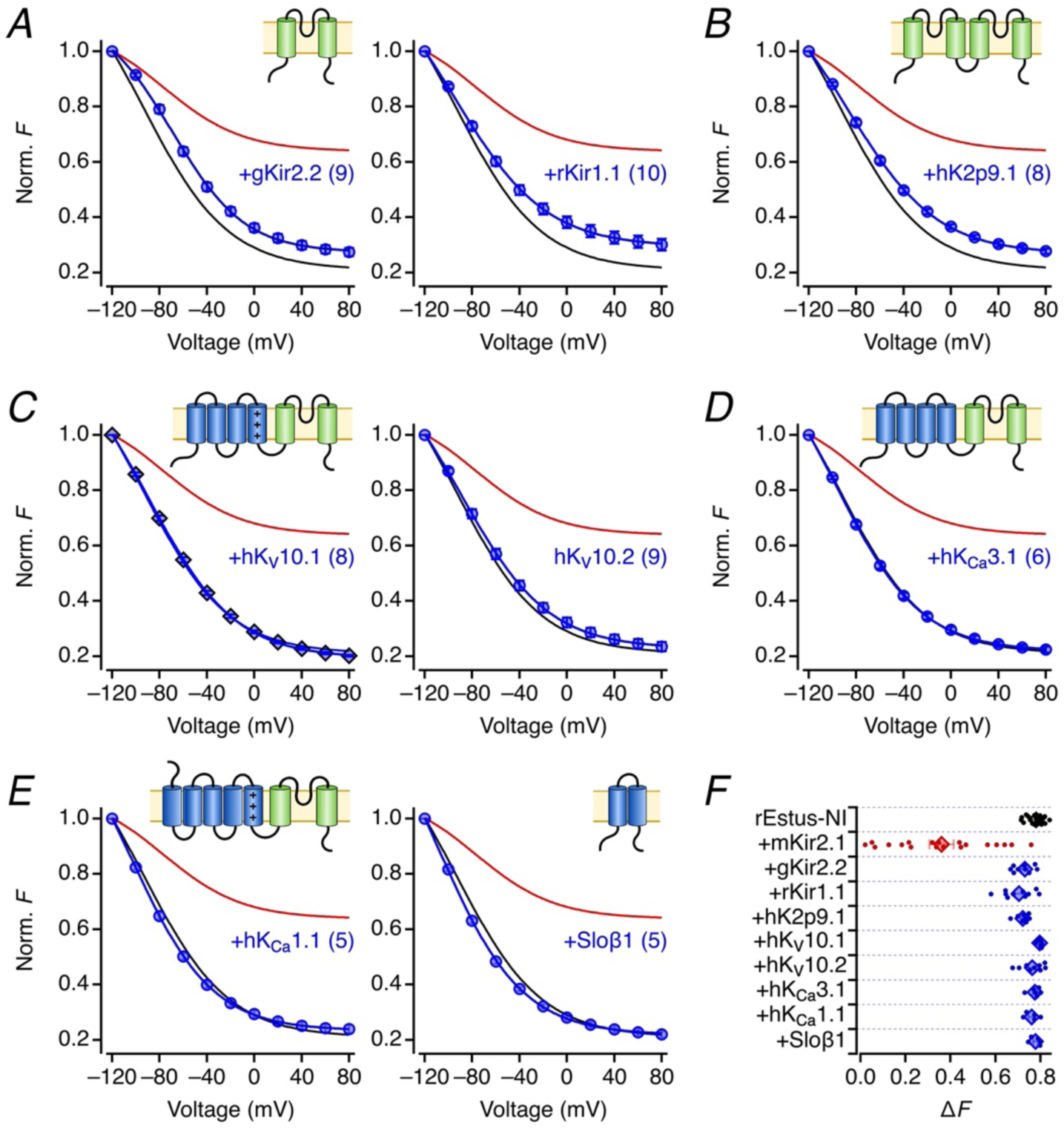
Fluorescence–voltage responses of rEstus-NI co-expressed with various K^+^ channels. *A*-*E*, Normalized *F*–*V* relationships of rEstus-NI in HEK293T cells recorded under whole-cell voltage clamp of rEstus-NI alone (black) and with co-expression of mKir2.1 (red) from Fig. 2A. The blue data points with superimposed Boltzmann fits are results from co-expression of the indicated channel constructs. The insets depict the membrane topology of the co-expressed proteins: (*A*) 2TM inward rectifier K^+^ channels gKir2.2 (IRK2, *kcnj12*) and rKir1.1 (ROMK1, *kcnj1*), (*B*) 2TM-2TM two-pore K^+^ channel K2p9.1 (TASK3, *KCNK9*), (*C*) 6TM voltage-gated K^+^ channels hKV10.1 (EAG1, *KCNH1*) and hKV10.2 (EAG2, *KCNH5*), (*D*) 6TM voltage-independent Ca^2+^-activated K^+^ channel hKCa3.1 (IKCa, *KCNN4*), and (*E*) 7TM Ca^2+^-activated K^+^ channels of large conductance hKCa1.1 (Slo1, BKCa, *KCNMA1*) or its 2TM β1 auxiliary subunit (Sloβ1, *KCNMB1*). Data are means ± SEM, with *n* in parentheses. Data for hKV10.1 were measured with 130 mM Cs^+^ and for hK2p9.1 with 14 mM K^+^ instead of 140 mM K^+^ as intracellular ion to limit the current size and, hence, to avoid series resistance problems. *F*, Normalized mean change in rEstus-NI fluorescence between -120 and 80 mV from the experiments in *A*-*E*.

Remarkably, co-expression of chicken Kir2.2 (gKir2.2) only marginally altered the *F*–*V* relationship of rEstus-NI, despite its close phylogenetic relationship to gKir2.1: Δ*F* 0.73 vs. 0.78 without gKir2.2 (*p* = 0.004, Fig. 4*A*). Similarly, the more distantly related rKir1.1 had only a minor effect on rEstus-NI voltage sensitivity (Fig. 4*A*): Δ*F* 0.70 vs. 0.78 without rKir1.1 (*p* = 6 10_– 5_, Fig. 4*A*). A similar result was obtained by co-expression of rEstus-NI with human K2p9.1 (TASK3), so-called two-pore K_+_ channels with a 2TM-2TM membrane topology: Δ*F* 0.72 vs. 0.78 without hK2p9.1 (*p* = 3 10_–4_, Fig. 4*B*). As for 2TM channels, hK2p9.1 had a weak impact on the performance of rEstus-NI. These results suggest that the observed strong alteration of the GEVI fluorescence response is a specific property of Kir2.1, but other inwardly rectifying K_+_ channels may also mildly interfere with rEstus-NI.

K_+_ channel subunits comprising their own functional VSD, such as the voltage-gated 6TM channels human K_V_10.1 (hK_V_10.1, EAG1) and K_V_10.2 (hK_V_10.2, EAG2), did not induce any change in the *F*–*V* of rEstus-NI (Fig. 4*C*). The α subunit of Ca^2+^-activated K^+^ channels hK_Ca_3.1 (intermediate-conductance, IK, *KCNN4*) also has a 6TM membrane topology, but its VSD is of minor functional relevance. When co-expressed with rEstus-NI, it did not affect the GEVIs voltage dependence either (Fig. 4*D*). α subunits of large-conductance Ca_2+_- and voltage-activated K^+^ channels of the BK type (hK_Ca_1.1, slo1) comprise seven transmembrane segments (7TM), with the additional S0 segment, rendering the N terminus on the extracellular side of the membrane, and a functional VSD formed by S1-S4. Neither co-expression of hK_Ca_1.1 with rEstus-NI nor even of its 2TM auxiliary β1 subunit (Sloβ1, *KCNMB1*) had a noticeable impact on rEstus-NI fluorescence (Fig. 4*E*). For the current responses during the voltage-clamp recordings of Fig. 4, refer to Supplementary Fig. 4; no obvious changes in the currents due to co-expression of rEstus-NI were noted.

In summary (Fig. 4*F*), these results indicate that the reduction of rEstus-NI voltage-dependent fluorescence responses observed with Kir2.1 is unique in its magnitude. There is a weak tendency for other channels with 2TM or 2TM-2TM motifs to have a minor impact on rEstus-NI function, but the K_+_ channel subunits with their own VSD do not share this property. Thus, it appears plausible that ASAP-type GEVIs form tetrameric complexes with Kir2.1 that resemble the architecture of 6TM Kv channels.

### VSDs interferes with Kir2.1 through a hydrophobic surface of their transmembrane domains

The results thus far presented indicate that mKir2.1 channels functionally interact with VSP-based GEVIs. We thus hypothesized that both protein subunits, consisting of a 4TM voltage-sensor motif and a 2TM pore-only motif, might form complexes that effectively constitute a non-covalently bound 6TM construct. We therefore used AlphaFold 3 to predict an octameric assembly consisting of four rEstus and four mKir2.1 subunits. The model with the highest-ranking score among five predicted models (ranking score 0.59, ipTM 0.50, pTM 0.53, no steric clashes detected) is shown in Fig. 5*A* and was selected for further analysis. The octameric protein complex was placed into a lipid membrane immersed in water with 150 mM KCl (Supplementary Fig. 5*A*). This simulation box was subjected to molecular dynamics simulation at 37°C for 100 ns. The stability of the structure (Supplementary Fig. 5*B*,*C*) indicates that the model is energetically plausible.

**Figure 5.**
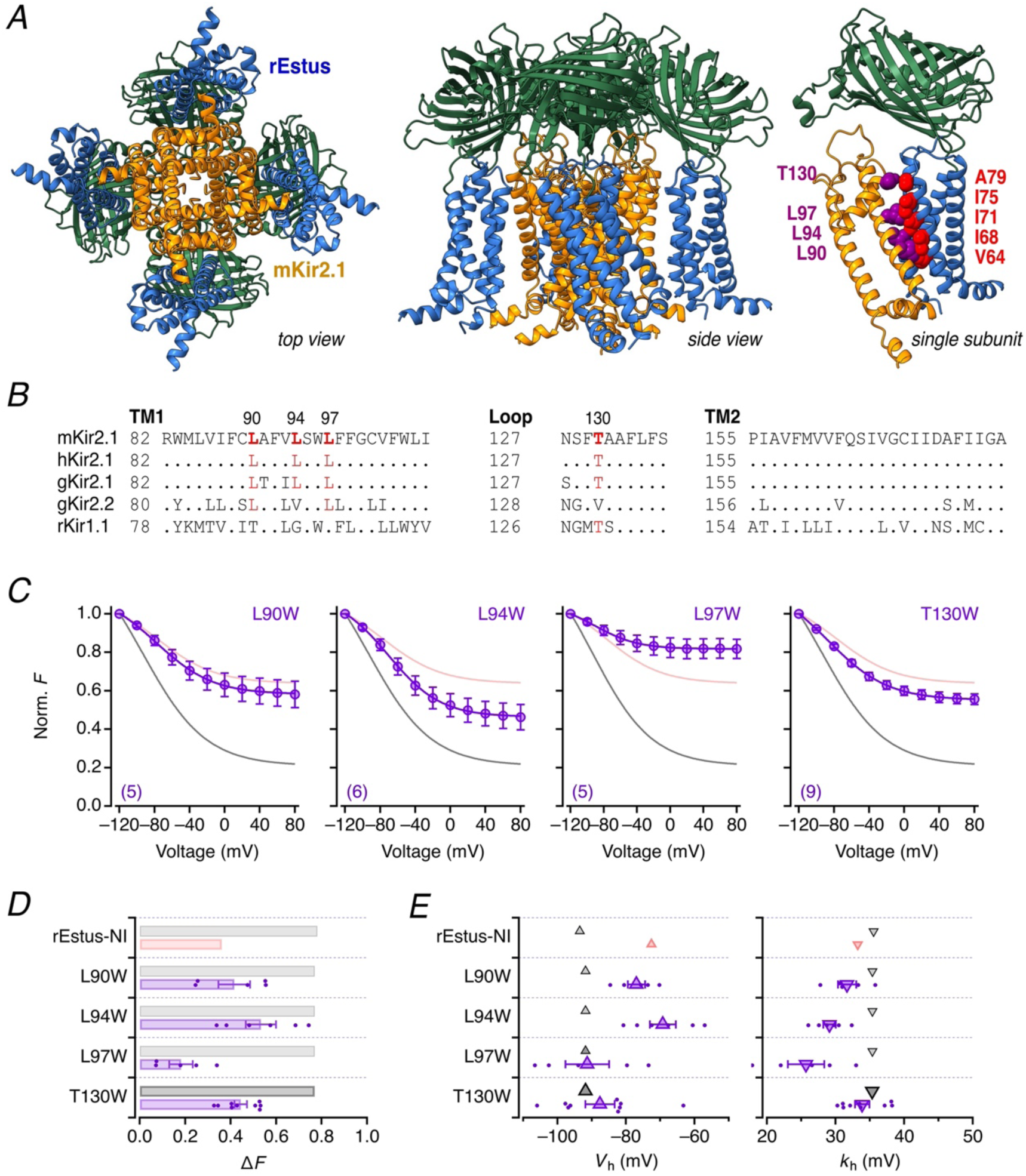
Structure prediction and experimental validation of mKir2.1 TM1 contribution. *A*, AlphaFold 3-predicted structural model of a protein complex consisting of four rEstus polypeptides co-assembled with a tetramer of mKir2.1 polypeptides shown in bottom view (*left*) and side view (*middle*). rEstus is shown in blue and mKir2.1 in yellow. The intracellular domains of mKir2.1 are not shown. *Right*: single-domain view highlighting residues in rEstus selected for mutagenesis (red, see Fig. 6) and nearby residues in mKir2.1 (purple). *B*, Multiple sequence alignment of the indicated Kir-channel types focusing on TM1, part of the loop between TM1 and the pore, and TM2. Residues identified in (*A*) as potentially interacting with rEstus-NI are shown in red. *C*, Normalized *F*–*V* relationships of rEstus-NI co-expressed with mKir2.1 variants mutated at predicted interface residues L90W, L94W, L97W, and T130W. Data are means ± SEM, with *n* in parentheses. The solid gray and pink curves are the reference fits for rEstus-NI and rEstus-NI plus mKir2.1, respectively. *D*, Mean relative change in fluorescence between -120 and 80 mV for rEstus-NI without (gray) or with co-expression of mKir2.1 (red) or mKir2.1 mutants (purple). *E*, Voltage of half-maximal GEVI activation (*V*h, *left*) and the corresponding slope factors (*k*h, *right*) from the data fits shown in (*C*). Mean data for rEstus-NI co-expressed with wild-type mKir2.1 are shown as reference.

This structural model shows tight packing between the first transmembrane helix of mKir2.1 (TM1) and S1 and S4 of the GEVI VSD (Fig. 5*A*, *right*) largely mediated by hydrophobic aliphatic side chains. L90, L94, and L97 in TM1, conserved in Kir2.1 from mouse, human, and chicken (Fig. 5*B*), are shown to interact with V64, I68, I71, and I75 in S1 of rEstus. In addition, A79 of rEstus is in close proximity to T130 in mKir2.1; the latter is part of the extracellular loop between TM1 and the selectivity filter, which is also conserved in mouse, human, and chicken Kir2.1 (Fig. 5*B*).

To assess the functional importance of the postulated tight packing, L90, L94, L97, and T130 were individually substituted with tryptophan, which has a rigid and bulky aromatic side chain. When co-expressed with rEstus-NI, these tryptophan-substituted mKir2.1 mutants retained the ability to alter the *F*–*V* relationship (Fig. 5*C-E*; *p* < 0.001 for all vs. rEstus-NI alone after correction for multiple comparisons). The effects of the L90W, L94W, L97W, and T130W mutants were not noticeably different from those of wild-type mKir2.1 (Fig. 5*D*). Thus, a single tryptophan substitution at the examined positions in mKir2.1 was insufficient to clearly disrupt the functional interaction between mKir2.1 and rEstus-NI.

The trend of an impaired functional interaction due to a mutation at position 94 in mKir2.1 (Fig. 5*C*) suggests that the non-conserved neighboring residues A91 and V93 might be causative for the observed difference between mKir2.1 and gKir2.1, the latter harboring the residues T91 and I93 (Fig. 5*B*). While currents mediated by the mKir2.1 mutants L90W, L94W, and L97W were similar to those of the wild type, mutant T130W yielded smaller currents (Supplementary Fig. 6).

To further evaluate the predicted interaction interface, the proposed interacting residues in rEstus-NI were also individually substituted with tryptophan. Without mKir2.1 co-expression, all single-site rEstus-NI tryptophan mutants possessed voltage-dependent fluorescence changes but the resulting *F*–*V* properties differed noticeably. The mutants rEstus-NI-I68W, I75W, and A79W retained *F*–*V* properties similar to the wild type (Fig. 6*A*, *D*, *E*). rEstus-NI-V64W and I71W displayed large left and right shifts in *V*_h_, respectively. For rEstus-NI-V64W, this left-shift was accompanied by a strongly increased *k*_h_, i.e., a lower steepness of the voltage dependence (Fig. 6*E*).

**Figure 6.**
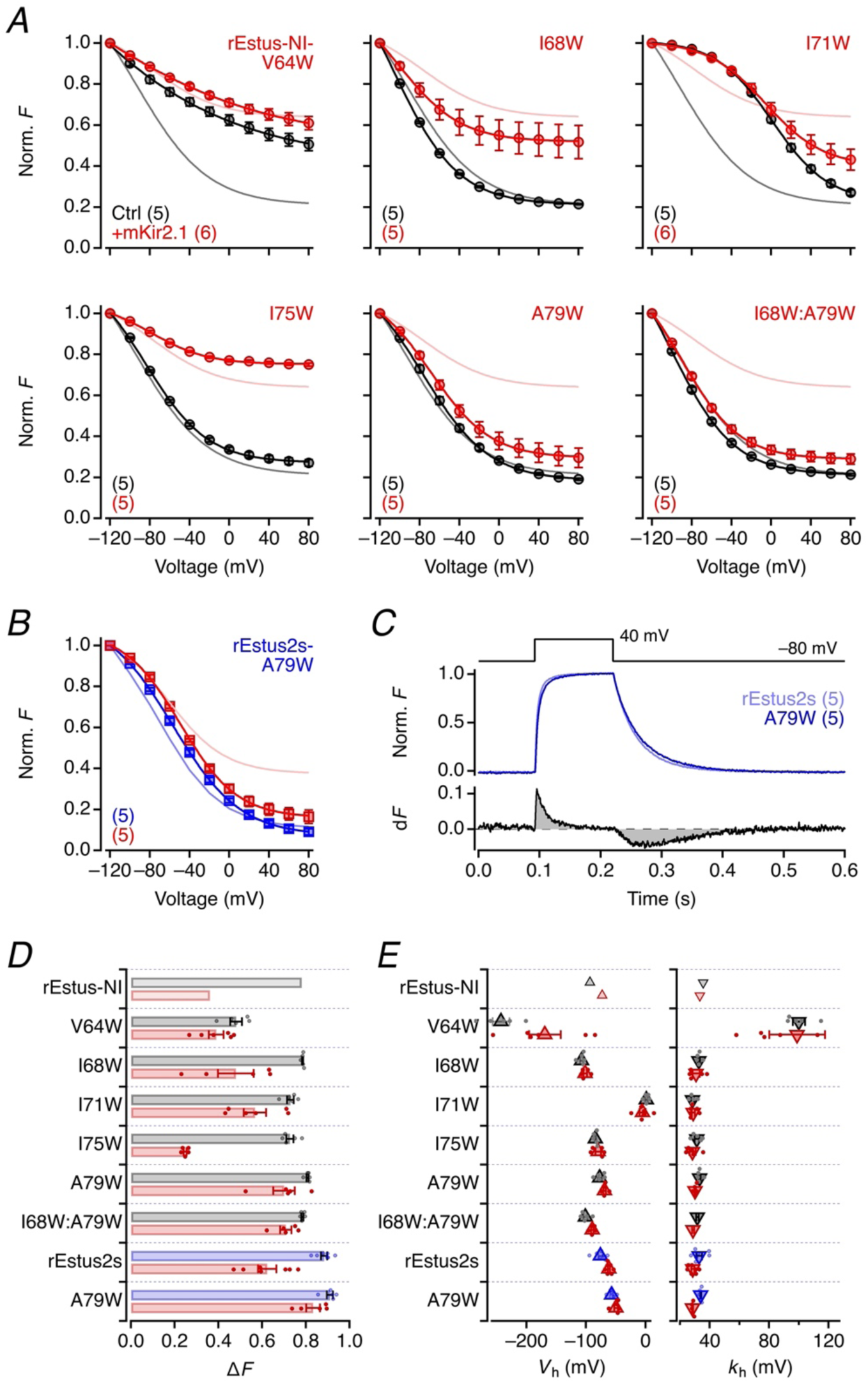
GEVI mutagenesis diminishes the impact of mKir2.1. *A*, Normalized fluorescence–voltage (*F*–*V*) relationships of rEstus-NI mutants carrying tryptophan substitutions at predicted interface residues (see Fig. 5A), expressed alone (black) or co-expressed with mKir2.1 (red). The last panel shows data for the double mutant rEstus-NI-I68W:A79W. Solid curves indicate Boltzmann fits (Eq. 1). Data are means ± SEM, with *n* in parentheses. Reference fits for rEstus-NI ± mKir2.1 (gray, pink). *B*, As in (*A*) for rEstus2s-A79W (blue) and co-expression with mKir2.1 (red). The fitted curves in light color are for rEstus2s without and with mKir2.1. *C*, Superposition of mean fluorescence recordings of rEstus2s (light blue) or rEstus2s-A79W (blue) for cells (*middle*), voltage-clamped according to the indicated protocol (*top*). The graph at the bottom shows the difference in normalized fluorescence between rEstus2s and rEstus2s-A79W to indicate a slightly slower response to the voltage steps, both in the depolarizing and the hyperpolarizing directions. *D*, Mean relative change in fluorescence between -120 and 80 mV for the indicated GEVIs and mutants without (rEstus-NI gray, rEstus2s blue) or with co-expression of mKir2.1 (red). *E*, Voltage of half-maximal GEVI activation (*V*h, *left*) and the corresponding slope factors (*k*h, *right*) from the fits to individual cell data as in (*A*) and (*B*).

Co-expression of mKir2.1 affected Δ*F* for rEstus-NI-I75W similarly to the wild type (from 0.73 ± 0.02 to 0.25 ± 0.01). For mutant I68W the impact of mKir2.1 on Δ*F* was diminished (from 0.76 ± 0.01 to 0.48 ± 0.08), while mutation A79W markedly diminished the impact of mKir2.1 on the *F*–*V* relationship (from 0.81 ± 0.01 to 0.70 ± 0.05). The I68W:A79W double mutation further reduced the impact of mKir2.1 (Δ*F* from 0.78 ± 0.01 to 0.71 ± 0.03). The markedly altered voltage dependence of mutants V64W and I71W precludes a direct comparison with wild-type rEstus-NI, but in both cases, co-expression of mKir2.1 had only a minor influence on the *F*–*V* relationship in the voltage range -120 to 80 mV (Fig. 6*A*, *D*). In summary, the I68W and A79W mutations substantially mitigated Kir2.1-induced attenuation of the GEVI response to *V*_m_ changes while maintaining near-native voltage sensitivity in the absence of mKir2.1 (Fig. 6*A*, *E*).

The rEstus-NI mutation A79W was selected for further analysis because it provided the most favorable balance between preserved voltage sensitivity and reduced susceptibility to Kir2.1-induced attenuation of Δ*F*. To determine whether this mutation effect generalized to other GEVIs, the A79W mutation was introduced in rEstus2s, a sensor with exceptional voltage sensitivity [30]. The mutation rEstus2s-A79W preserved the overall sensor functions with only a minor slowing of the fluorescence response in the depolarizing and the hyperpolarizing directions, compared with rEstus2s alone (Fig. 6*C*). However, the mutation virtually eliminated the influence of mKir2.1 co-expression (Fig. 6*B*, *D*).

## Discussion

We have demonstrated that co-expression of Kir2.1 channels of the 2TM family, comprising only a pore domain (PD), markedly alters the fluorescence signal from ASAP-type GEVIs such as rEstus-NI derived from a voltage-sensitive phosphatase. Kir2.1 attenuates the maximal voltage-dependent fluorescence signal and reshapes the fluorescence-voltage relationship. The magnitude of the fluorescence change varied among cells, suggesting that the interaction depends on the relative expression levels of Kir2.1 and GEVI. The observed functional interaction is highly specific to the Kir2.1 channel. While mouse, human, and chicken Kir2.1 all alter the GEVI fluorescence signal, the closely related Kir2.2, the more distantly related Kir1.1, and other channels including two-pore K_+_ channels are much less effective.

The functional interaction between Kir2.1 channels of the 2TM family and ASAP-type GEVIs with VSDs can be interpreted in the context of the overall structure of tetrameric Kv channel complexes, in which each subunit contains one VSD and one PD (Fig. 1*B*). For example, in Kv1 channels, the VSD and PD are tightly packed through interactions between S1 and S5 (the latter being equivalent to TM1 in Kir2.1) and the two segments surround S4 with positive voltage-sensing charges. Given the aforementioned structural design and the well-known modular assembly of proteins including ion channels, as well illustrated in the “split channel” experiments [12, 13], it is perhaps mechanistically plausible that two separate proteins, Kir2.1 and rEstus-NI, each of which is functional on its own, may assemble into a non-covalent Kv-like architecture with altered functional properties in which the Kir2.1 PD substitutes for the S5-S6 PD of a conventional Kv subunit. This may mirror the evolutionary emergence of Kv channels because 2TM channels and VSD-only proteins, such as VSPs and proton channels lacking a PD, are considered to predate Kv channels. This notion is supported by our observation that gKir2.1 has a stronger impact on rEstus-NI, with its VSD from chicken, than human Kir2.1. This may indicate a result of evolution with a potential functional assembly of VSPs and Kir2.1 channels.

A structural model with four Kir2.1 subunits and four rEstus subunits generated with AlphaFold 3 and refined by molecular dynamics simulations shows a putative physical interaction interface largely mediated by hydrophobic aliphatic side chains between Kir2.1 and rEstus. Guided by this model, mutagenesis indeed shows that select mutations in mKir2.1 TM1 and GEVI S1 disrupt the functional interaction. While the results presented alone do not conclusively show that Kir2.1 and ASAP-type GEVI indeed assemble into a Kv-type complex, the agreement between the structural model and the mutagenesis results strengthens the plausibility of the proposed non-covalent Kv-like assembly where the VSDs and the PD are positioned for functional coupling. The coupling mechanism between the Kir2.1 PD and rEstus- NI VSDs could involve PI(4,5)P₂ (e.g., [35]), as Kir2.1 is considered to be lipid-gated channels.

In ASAP-type GEVIs, the fluorescence excited at 480 nm originates predominantly from the deprotonated anionic chromophore and is highly voltage sensitive, whereas fluorescence excited at 400 nm from the protonated phenolic chromophore is much less voltage dependent (e.g., [21] and Supplementary Fig. 2*B*). The relative population between the two chromophore states is indirectly influenced by the state of the VSD and its local environment. Operationally, the fluorescence ratio *F*_480_/*F*_400_ (*r*) provides an absolute estimate of the sensor configuration. This framework can be used to interpret the changes in the *F*–*V* relationship produced by Kir2.1. In rEstus-NI, mKir2.1 co-expression restricted the dynamic range of the fluorescence ratio: *r*_max_ at negative voltages decreased, whereas *r*_min_ at positive voltages increased, resulting in a crossover of the control and Kir2.1 curves near −60 mV (Fig. 3*D*). One explanation is that interaction with Kir2.1 alters the relative occupancies of fluorescence-reporting states of rEstus-NI, reflecting differences in VSD configuration, chromophore states and/or their coupling. Trapping of VSD in an intermediate state has been demonstrated in Ci-VSP where the VSD may be prevented from undergoing the full range of voltage-dependent conformational changes and therefore never reach the conformation associated with the brightest and dimmest levels of fluorescence [36]. Unlike rEstus-NI, ASAP5 exhibited a distinct response to Kir2.1 co-expression: *r*_max_ at negative voltages decreased and *r*_min_ at positive voltages remained essentially unaltered. The results together indicate that Kir2.1 differentially alters the relative occupancy of fluorescence-reporting states in different ASAP-type GEVIs. One possible contributor to the observed differences is the linker connecting the cpEGFP to the VSD. ASAP5 and JEDI-1P contain the linker sequences LAAFNSH and LAAFNSE, respectively, whereas rEstus-NI, rEstus, ASAP3, and rEstus2s share the GTFRGD linker. Differences in linker length (7 vs. 6 residues) or sequence could alter the coupling between VSD conformational changes and the chromophore, thereby reducing the functional consequences of Kir2.1 co-expression.

The experimental findings are generally consistent with the proposed non-covalent Kv-like assembly; four Kir2.1 and four ASAP-type subunits, such as rEstus-NI, form an octameric complex. However, additional lines of evidence are clearly required to substantiate the model. Biochemical approaches including direct structural determinations will be particularly informative. Further, some important questions remain unresolved. While the structural model assumes a 4:4 stoichiometry of Kir and rEstus in one functional complex, the exact stoichiometry is unknown and it may not be fixed. Furthermore, it is not clear if the VSD engages with Kir proteins as monomers or as dimers. It will be important to determine when the Kir2.1-rEstus complex forms during biogenesis, for example, in the endoplasmic reticulum or after trafficking to the plasma membrane. Yet another unresolved question concerns the functional properties of the channel Kir2.1. Our study clearly shows that the fluorescence properties of rEstus-NI are influenced by Kir2.1. But allosteric interactions within proteins are generally reciprocal in theory and some effects on Kir2.1 might be expected. However, our electrophysiological measurements do not reveal any obvious effects on ionic currents mediated by Kir2.1 by rEstus-NI. Despite these open questions, an interaction of Kir2.1 with other membrane proteins is not unprecedented. For example, Kir2.1 participates in macromolecular complexes with cardiac voltage-gated sodium channels (Na_V_1.5), with which it forms a cardiac “channelosome” that reciprocally shapes excitability [37, 38]. In this proposed interaction, S1 of Na_V_1.5’s VSD of domain II also engages TM1 of Kir2.1 [38]. Moreover, hydrophobic residues in S1 are essential for the voltage-sensor function in voltage-sensitive phosphatases [39], which is compatible with our results that S1 mutations in rEstus-NI (Fig. 6) and an interference of Kir2.1 TM1 with S1 in an ASAP-type GEVI can affect its voltage sensitivity.

Our findings here have clear practical and cautionary implications for the use of ASAP-type GEVIs in cell physiological experiments. Under conditions where GEVI expression is low or Kir2.1 expression is high, such as in cardiomyocytes, ASAP-type GEVIs may exhibit reduced signal amplitude. Since Kir2.1 is also commonly used during GEVI screening, it may inadvertently influence the development of new sensors. For precise determination of *V*_m_ using GEVIs, this interaction may still pose a technical challenge if Kir2.1 channel expression varies among cells used for sensor calibration and those used in the final experimental setting. Therefore, GEVIs that are less susceptible to Kir channel interference will be valuable. The A79W mutation, shown here for rEstus-NI and rEstus2s (Fig. 6), mitigates this effect and may be incorporated into future GEVI designs to prevent this potential artifact. Kir2.1 may represent only one example of a broader class of unintended interactions between GEVIs and other membrane proteins. Such unforeseen interactions may alter and degrade sensor performance in a cell type-dependent fashion. This issue should always be considered during sensor development and experimental design.

Our findings also imply that the functional interaction of Kir2.1 with unrelated, VSD-only protein family members may be a more general property of Kir2.1 than previously appreciated.

## Additional information

### Data availability statement

All data supporting the results are presented in the published paper or the supplementary material.

### Author contributions

A.G.N. data collection, analysis, writing; P.R. study design, generation of mutants, structural modeling, instruction; L.S. data collection; T.H. molecular dynamics simulation; R.S. biochemical assays; S.H.H. study design, data analysis, figure generation, writing, fund raising. All authors contributed to the final editing of the text.

## References

1. Doyle, D.A., et al., The structure of the potassium channel: molecular basis of K^+^ conduction and selectivity. Science, 1998. 280(5360): p. 69–77.

2. MacKinnon, R., Pore loops: an emerging theme in ion channel structure. Neuron, 1995. 14(5): p. 889–92.

3. Thiel, G., et al., Potassium ion channels: could they have evolved from viruses? Plant Physiology, 2013. 162(3): p. 1215–1224.

4. Vardanyan, V. and O. Pongs, Coupling of voltage-sensors to the channel pore: a comparative view. Front Pharmacol, 2012. 3: p. 145.

5. Heiss, M.C. and B.E. Flucher, Voltage-sensing domains: structural and functional diversity. Eur Biophys J, 2025.

6. DeCoursey, T.E., Voltage and pH sensing by the voltage-gated proton channel, HV1. J R Soc Interface, 2018. 15(141).

7. Ramsey, I.S., et al., A voltage-gated proton-selective channel lacking the pore domain. Nature, 2006. 440(7088): p. 1213–1216.

8. Sasaki, M., M. Takagi, and Y. Okamura, A voltage sensor-domain protein is a voltage-gated proton channel. Science, 2006. 312(5773): p. 589–92.

9. Matsuda, M., et al., Crystal structure of the cytoplasmic phosphatase and tensin homolog (PTEN)-like region of Ciona intestinalis voltage-sensing phosphatase provides insight into substrate specificity and redox regulation of the phosphoinositide phosphatase activity. Journal of Biological Chemistry, 2011. 286(26): p. 23368–23377.

10. Murata, Y., et al., Phosphoinositide phosphatase activity coupled to an intrinsic voltage sensor. Nature, 2005. 435(7046): p. 1239–1243.

11. St-Pierre, F., et al., High-fidelity optical reporting of neuronal electrical activity with an ultrafast fluorescent voltage sensor. Nature Neuroscience, 2014. 17(6): p. 884–889.

12. Lörinczi, É., et al., Voltage-dependent gating of KCNH potassium channels lacking a covalent link between voltage-sensing and pore domains. Nat Commun, 2015. 6: p. 6672.

13. de la Peña, P., P. Domínguez, and F. Barros, Functional characterization of Kv11.1 (hERG) potassium channels split in the voltage-sensing domain. Pflugers Arch, 2018. 470(7): p. 1069–1085.

14. Hibino, H., et al., Inwardly rectifying potassium channels: their structure, function, and physiological roles. Physiol Rev, 2010. 90(1): p. 291–366.

15. de Boer, T.P., et al., The mammalian K(IR)2.x inward rectifier ion channel family: expression pattern and pathophysiology. Acta Physiol (Oxf), 2010. 199(3): p. 243–56.

16. Dhamoon, A.S. and J. Jalife, The inward rectifier current (IK1) controls cardiac excitability and is involved in arrhythmogenesis. Heart Rhythm, 2005. 2(3): p. 316–324.

17. Sakmann, B. and G. Trube, Voltage-dependent inactivation of inward-rectifying single-channel currents in the guinea-pig heart cell membrane. J Physiol, 1984. 347: p. 659–83.

18. Schwarz, W., B. Neumcke, and P.T. Palade, K-current fluctuations in inward-rectifying channels of frog skeletal muscle. J Membr Biol, 1981. 63(1-2): p. 85–92.

19. Nakajima, Y., S. Nakajima, and M. Inoue, Pertussis toxin-insensitive G protein mediates substance P-induced inhibition of potassium channels in brain neurons. Proceedings of the National Academy of Sciences, 1988. 85(10): p. 3643–3647.

20. Brismar, T. and V.P. Collins, Inward rectifying potassium channels in human malignant glioma cells. Brain Res, 1989. 480(1-2): p. 249–58.

21. Rühl, P., et al., An ultrasensitive genetically encoded voltage indicator uncovers the electrical activity of non-excitable cells. Advanced Science, 2024. 11(20).

22. Abramson, J., et al., Accurate structure prediction of biomolecular interactions with AlphaFold 3. Nature, 2024. 630(8016): p. 493–500.

23. Pettersen, E.F., et al., UCSF ChimeraX: Structure visualization for researchers, educators, and developers. Protein Sci, 2021. 30(1): p. 70–82.

24. Wu, E.L., et al., CHARMM-GUI Membrane Builder toward realistic biological membrane simulations. J Comput Chem, 2014. 35(27): p. 1997–2004.

25. van Meer, G., D.R. Voelker, and G.W. Feigenson, Membrane lipids: where they are and how they behave. Nat Rev Mol Cell Biol, 2008. 9(2): p. 112–24.

26. Nair, A.G., et al., Absolute membrane potential recording with ASAP-type genetically encoded voltage indicators using fluorescence lifetime imaging. ACS Chemical Neuroscience, 2025. 16(24): p. 4636–4646.

27. Villette, V., et al., Ultrafast two-photon imaging of a high-gain voltage indicator in awake behaving mice. Cell, 2019. 179(7): p. 1590–1608.e23.

28. Lu, X., et al., Widefield imaging of rapid pan-cortical voltage dynamics with an indicator evolved for one-photon microscopy. Nature Communications, 2023. 14(1): p. 6423–6423.

29. Hao, Y.A., et al., A fast and responsive voltage indicator with enhanced sensitivity for unitary synaptic events. Neuron, 2024. 112(22): p. 3680–3696.e8.

30. Rühl, P., et al., Sub-millivolt voltage imaging reveals gap junction-mediated bioelectric contact inhibition. Nature Communications, 2026. in press.

31. Madeira, F., et al., The EMBL-EBI Job Dispatcher sequence analysis tools framework in 2024. Nucleic Acids Res, 2024. 52(W1): p. W521–w525.

32. Schindelin, J., et al., Fiji: an open-source platform for biological-image analysis. Nat Methods, 2012. 9(7): p. 676–82.

33. Rühl, P., et al., Monitoring of compound resting membrane potentials of cell cultures with ratiometric genetically encoded voltage indicators. Commun Biol, 2021. 4(1): p. 1164.

34. Ma, D., et al., An andersen-Tawil syndrome mutation in Kir2.1 (V302M) alters the G-loop cytoplasmic K^+^ conduction pathway. J Biol Chem, 2007. 282(8): p. 5781–9.

35. Hansen, S.B., Lipid agonism: The PIP2 paradigm of ligand-gated ion channels. Biochim Biophys Acta, 2015. 1851(5): p. 620–8.

36. Shen, R., et al., Mechanism of voltage gating in the voltage-sensing phosphatase Ci-VSP. Proc Natl Acad Sci U S A, 2022. 119(44): p. e2206649119.

37. Gutiérrez, L.K., A.I. Moreno-Manuel, and J. Jalife, Kir2.1-NaV1.5 channelosome and its role in arrhythmias in inheritable cardiac diseases. Heart Rhythm, 2024. 21(5): p. 630–646.

38. Stary-Weinzinger, A., In silico models of the macromolecular NaV1.5-KIR2.1 complex. Front Physiol, 2024. 15: p. 1362964.

39. Rayaprolu, V., et al., Hydrophobic residues in S1 modulate enzymatic function and voltage sensing in voltage-sensing phosphatase. J Gen Physiol, 2024. 156(7).

