## Supplementary Information for "Structural compatibility enables functional co-assembly of Kir2.1 channels and the voltage-sensing domain of voltage-sensitive phosphatases"

Anagha Gopalakrishnan Nair[1#], Philipp Rühl[1#], Lina Seeber[1], Toshinori Hoshi[2], Roland Schönherr[1], Stefan H. Heinemann[1*]

### Shared first authorship.

1 Center for Molecular Biomedicine, Department of Biophysics, Friedrich Schiller University Jena and Jena University Hospital, Jena, Germany.

2 Department of Physiology, University of Pennsylvania, Philadelphia, USA.

*Corresponding author:

Center for Molecular Biomedicine, Department of Biophysics, Friedrich Schiller University Jena & Jena University Hospital, Hans-Knöll-Str. 2, 07745 Jena, Germany.

#### Supplementary Figures

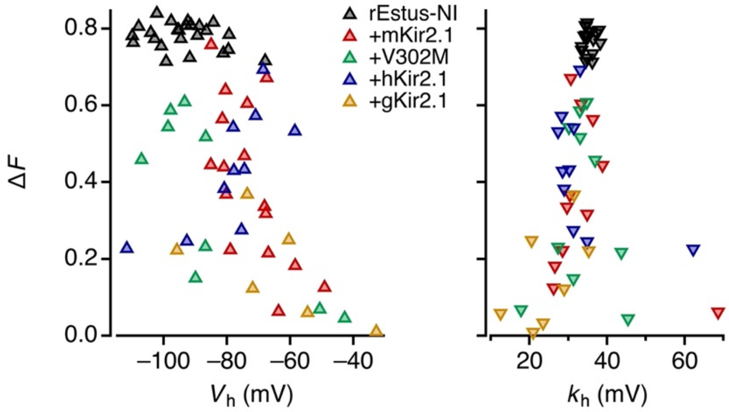


**Supplementary Figure 1. Correlation of parameters describing the voltage dependence with the maximal rEstus-NI fluorescence change**

Scatter plots of the data shown in Fig. 2 for rEstus-NI alone (black) or co-expressed with one of the indicated Kir2.1 variants (colored). The half-maximal fluorescence change voltage ($V_h$) tended to move toward more depolarized $V_h$ values as $\Delta F$ decreased (*left*), while the slope factor $k_h$ tended to decrease with smaller $\Delta F$ values (*right*).

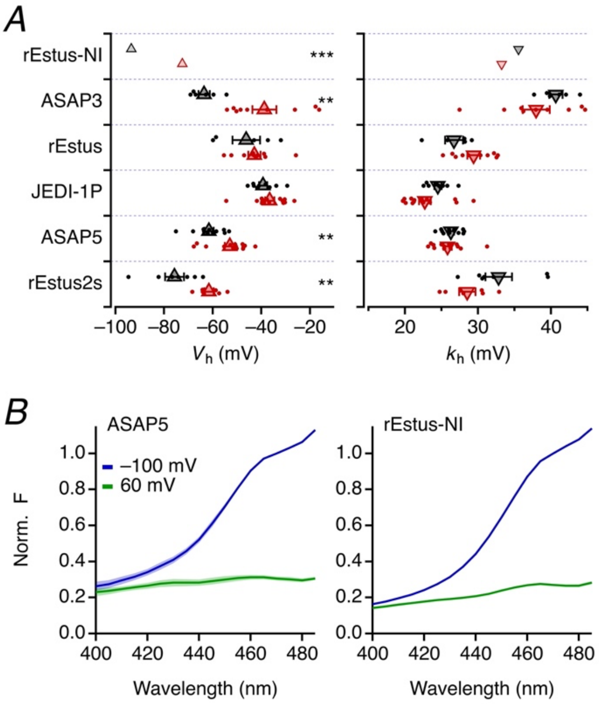


**Supplementary Figure 2. Functional GEVI properties (supplement to Figure 3)**

*A*, Voltage of half-maximal GEVI activation ($V_h$, *left*) and the corresponding slope factors ($k_h$, *right*) from the data fits shown in Figure 3*A*; mean values for rEstus-NI are shown as reference. *B*, Excitation spectra, normalized to 470 nm, of ASAP5 and rEstus-NI in HEK293T cells at -100 and 60 mV. Data are means ± sem in shading, *n* = 5. The spectra are corrected for wavelength-dependent excitation light intensity variations.

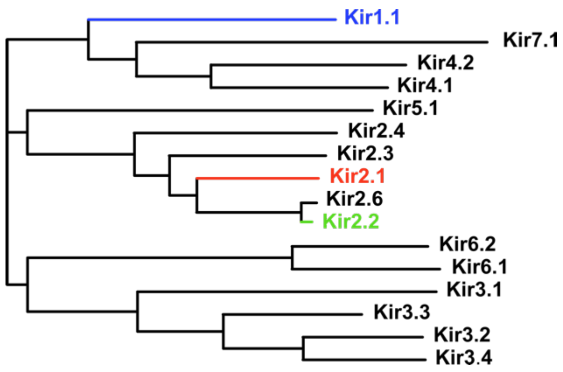


**Supplementary Figure 3. Phylogenetic tree of inward rectifier channel subunits**

Protein sequences of human inward rectifier channel subunits were retrieved from UniProt and aligned using the UniProt multiple sequence alignment tool. The resulting phylogenetic tree was visualized using iTOL (Interactive Tree of Life: itol.embl.de). The colored variants were examined in this study. Alignments of the TM1 and TM2 transmembrane regions revealed an identity between human Kir2.1 and Kir2.2 of 53.6 and 67.9%, respectively, while these scores were 40 and 54.4% for Kir2.1 compared with Kir1.1.

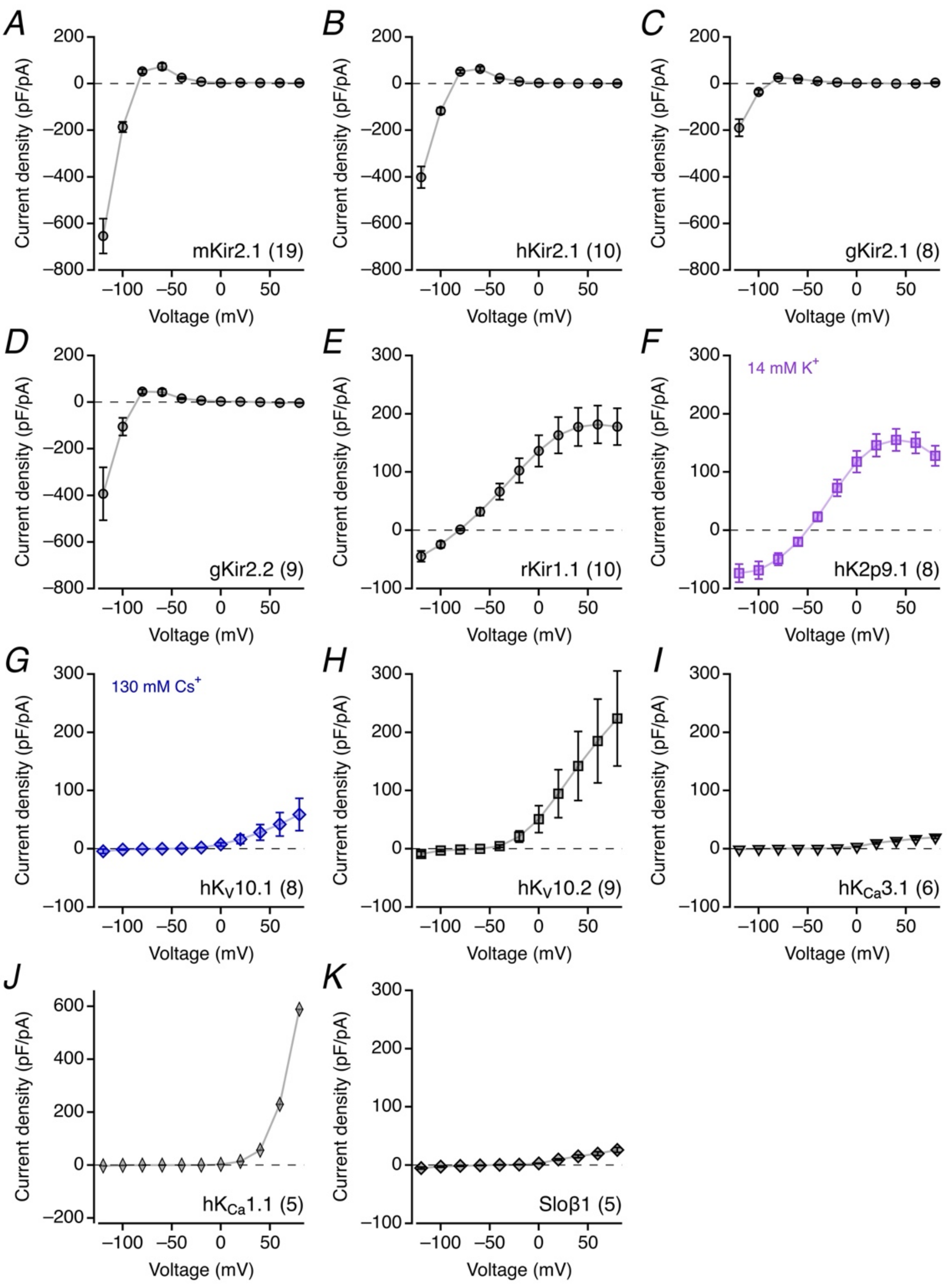


**Supplementary Figure 4. Current recordings from HEK293T cells expressing rEstus-NI along with various $K^+$ channel types**

*A-K*, Mean current densities (mean current during the second half of the depolarization segment (see Fig. 2*B*) divided by the cell capacitance) as a function of voltage from HEK293T cells expressing rEstus-NI along with the indicated $K^+$ channel types. Data are means ± SEM, *n* in parentheses. Straight lines connect the data points. Because very large currents distort the real applied voltage due to limited series resistance compensation, currents for hK2p9.1 (*F*) and $hK_V10.1$ (*G*) were measured with pipette solutions containing 14 mM $K^+$ 130 mM $Cs^+$ instead of 140 mM $K^+$ ions. Note that the intracellular solutions were buffered with 10 mM EGTA, thus eliminating most of the free $Ca^{2+}$ ions. Therefore, $hK_{Ca}3.1$ (*I*) does not show current above the leak level and $hK_{Ca}1.1$ only starts activating above about 40 mV. In the presence of the β1 subunit (*K*), activation of $hK_{Ca}1.1$ is right-shifted on the voltage axis and, thus, also does not produce $K^+$ currents in the voltage range examined.

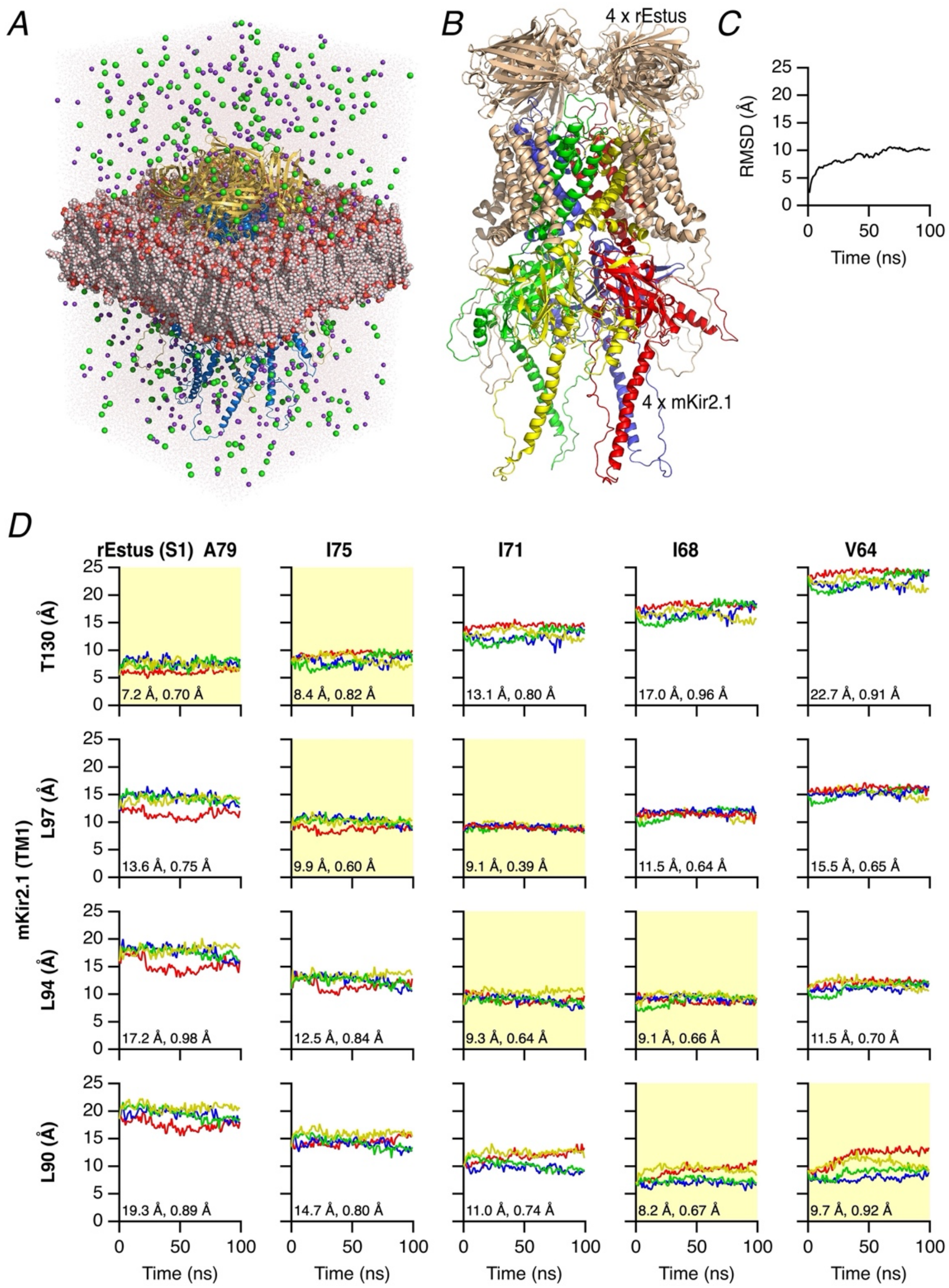


**Supplementary Figure 5. Molecular dynamics simulation of an octameric rEstus–mKir2.1 assembly in a lipid membrane**

*A*, Simulation box consisting of four rEstus polypeptides, four mKir2.1 polypeptides, a lipid membrane, water, and 150 mM KCl. *B*, Side view of the octameric rEstus–mKir2.1 complex. rEstus is shown in gold, the four units of mKir2.1 are colored. *C*, Root mean squared deviation (RMSD) of all protein atoms during the simulation of 100 ns indicates that the complex reaches an equilibrium structure after about 60 ns. *D*, Distances between the alpha carbon (CA) atoms of the indicated residues in mKir2.1 and the voltage-sensor domain of rEstus as a function of simulation time. Results from the four subunits are indicated in colors matching the color-coding of panel (*B*). The numbers in the panels indicate the mean distance in all four subunits and the mean SEM of the simulation time courses. Yellow shading indicates residue pairs with the two lowest mean distances. Lowest SEM values indicate a firm interaction between the respective amino acid residues: T130-A79, L97-I75, L94-I71, L90-I68.

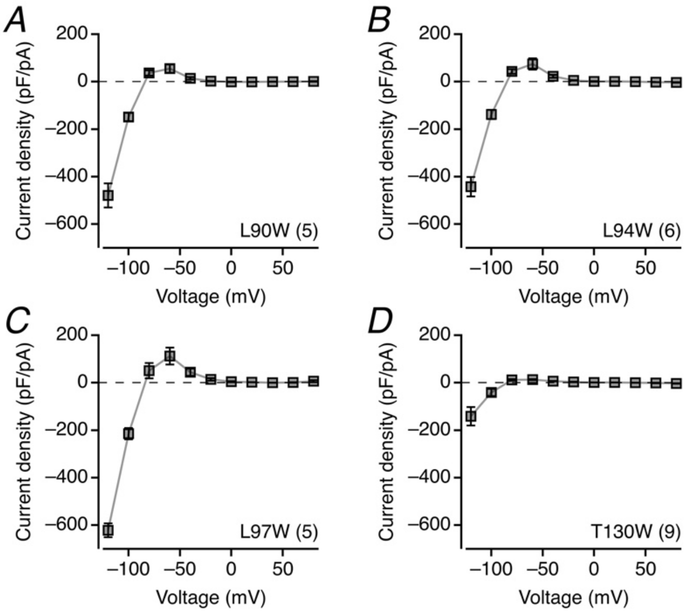


**Supplementary Figure 6. Current recordings from HEK293T cells expressing rEstus-NI along with mKir2.1 mutants**

*A*-*D*, Average current densities (mean current during the second half of the depolarization segment (see Fig. 2*B*) divided by the cell capacitance) as a function of voltage from HEK293T cells expressing rEstus-NI along with the indicated mutants of mKir2.1. Data are means ± SEM, *n* in parentheses. Straight lines connect the data points.
